# Design Space Exploration of the Violacein Pathway in *Escherichia coli* Based Cell-Free System

**DOI:** 10.1101/027656

**Authors:** Phuc H.B. Nguyen, Yong Y. Wu, Shaobin Guo, Richard M. Murray

## Abstract

In this study, an *Escherichia coli* (*E. coli*) based transcription translation cell-free system (TX-TL) was employed to sample various enzyme expression levels of the violacein pathway. The pathway was successfully reconstructed in TX-TL. Its variation produced different metabolites as evident from the extracts’ assorted colors. Analysis of the violacein product via UV-Vis absorption and liquid chromatography-mass spectrometry (LC-MS) detected 68 ng of violacein per μl reaction volume. Significant buildup of prodeoxyviolacein intermediate was also detected in the equimolar TX-TL reaction. Finally, design space exploration experiments suggested an improvement in violacein production at high VioC and VioD DNA concentrations.

## Introduction

Metabolic engineering carries the potential of inexpensive and simplified synthesis of complex biological molecules that would otherwise be too expensive to produce. The field has made important contributions to the industrial synthesis of many biomolecules such as ethanol, glycerol, and lysine [1]. However, many engineered metabolic pathways are poorly optimized, leading to buildup of toxic intermediates and byproducts and limiting cell growth. This in turn resulted lowered yield and efficiency [2]. To resolve this problem, multiple cycles of design iteration of the synthetic pathway are performed until an optimized version is obtained. Nevertheless, this process is time-consuming and inefficient. Cloning and transforming multiples genes in bacteria can be labor-intensive and take at least 1 week for each iteration [3]. This inevitably lengthens the cycle time required for testing and data collecting.

In this paper we make use of an E. coli-based, cell-free transcription-translation (TX-TL) system that employs both linear and circular DNA. TX-TL enables a fast and simple in vitro testing platform for biological genetic networks [4]. First, DNA parts of interest are created using PCR. These pieces are ligated together using Golden Gate cloning method. Complete assembly constructs include promoter, UTR, CDS, and terminator. Finally, these constructs are tested *in vitro*. This enables a time-efficient alternative for the conventional *in vivo* expression method. Instead of going through the lengthy transformation and bacterial culturing process, enzyme’s DNA sequences can simply be assembled into different linear DNAs and expressed directly in TX-TL. This significantly reduces the time required for testing of each expression variation and allow for a more efficient exploration of metabolic design space.

Violacein is a water-insoluble bacterial pigment with potential applications such as antibacterial, anti-trypanocidal, anti-ulcerogenic, and anticancer drugs [5]. The compound is natively synthesized by *Chromobacterium violaceum* [6]. Nevertheless, its production is too expensive due to the strain’s low productivity [5]. The violacein production pathway is shown in Fig. 1. The five core enzymes vioA-E of the pathway have been well-defined [6]. The starting source of violacein synthesis is tryptophan. The amino acid is then processed by a series of enzymes called vioA-E and turned into the following intermediates, respectively, before being converted into violacein: indole-3-pyruvate, X, prodeoxyviolacein, and proviolacein. Here, X represents an unknown intermediate. Without the presence of vioE, it can automatically be converted into chromopyrrolic acid. A common contaminant of the pathway, deoxyviolacein, can be created if prodeoxyviolacein is processed by vioC instead of vioD.

**Fig. 1.**
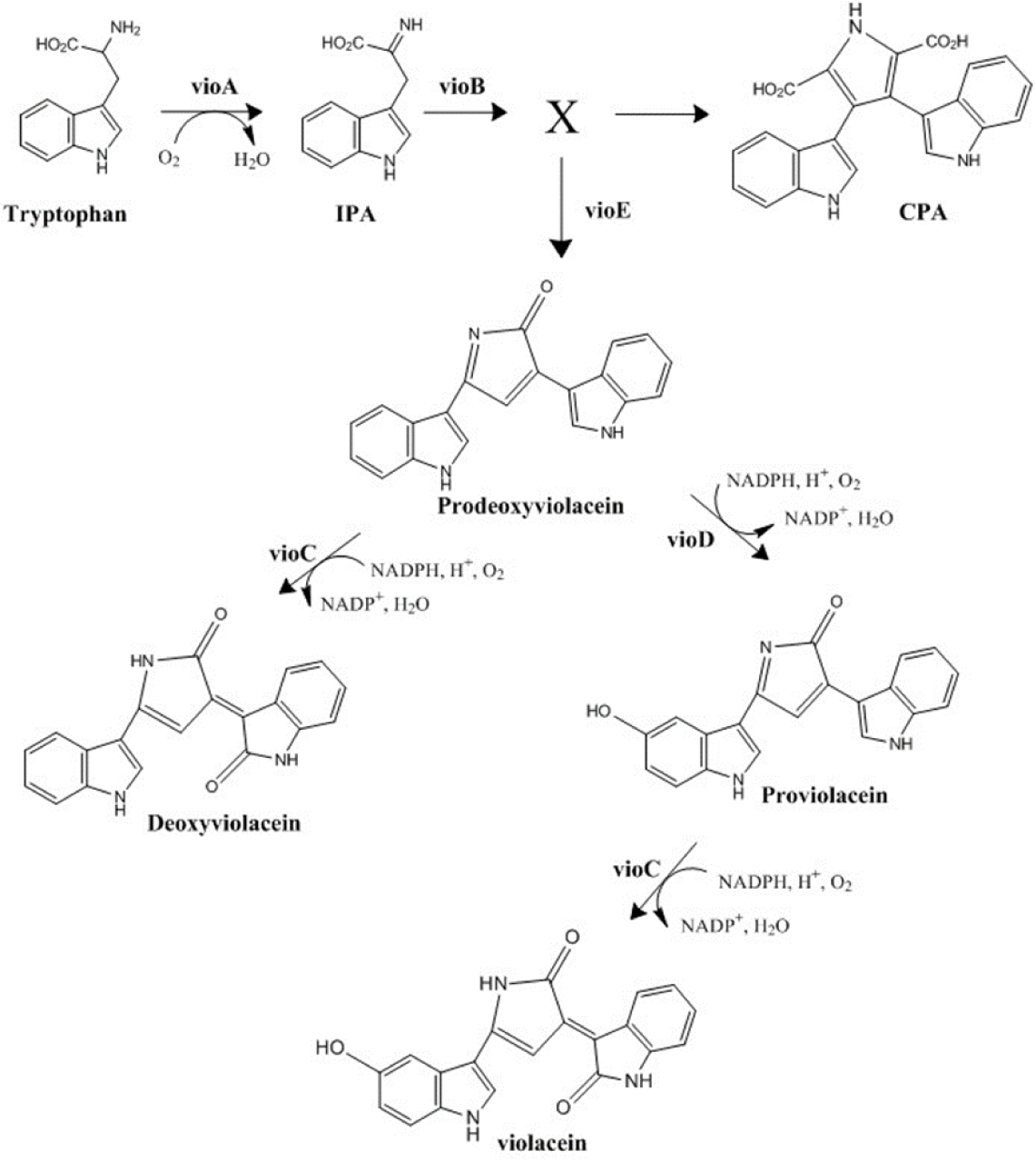
Overview of violacein production pathway. The starting source of violacein synthesis is tryptophan. The amino acid is then processed by a series of enzymes called VioA-E and turned into the following intermediates, respectively, before being converted into violacein: indole-3-pyruvate, X, prodeoxyviolacein, and proviolacein. Here, X represents an unknown intermediate. Without the presence of VioE, it can automatically be converted into chromopyrrolic acid. A common contaminant of the pathway, deoxyviolacein, can be created if prodeoxyviolacein is processed by VioC instead of VioD.

We seek to employ TX-TL to rapidly prototype the violacein pathway for improved product yield. By testing different expression levels of violacein pathway enzymes, valuable information such as rate-limiting step as well as the pathway’s dynamic can be obtained. Here, we successfully reconstructed the violacein pathway in TX-TL. We demonstrated that the CDS length of each enzyme significantly affects its expression level. Finally, prototyping results suggest a high VioC and VioD DNA concentrations results in an improved violacein production while minimizing intermediate buildup. These results can be used for the future engineering of more efficient violacein producing *E.coli* strain.

## Results

### TX-TL reactions of violacein pathway and its variants yielded various colored products

Coding sequences (CDS) of VioA-E enzymes [2] were cloned under the same promoter and RBS to minimize variation in expression levels at the same DNA concentration. Here, the violacein pathway and its modifications were reconstructed by simply leaving out specific enzymes DNA in TX-TL. The reactions were run using equimolar amount of each vio enzyme’s linear DNA. The products were found to be insoluble and extracted using 70% ethanol. The result indicated production of different metabolites as evident from the extracts’ various colors (Fig. 2). The extract from the complete pathway producing violacein had a deep violet color as expected. Extract from dexoyviolacein pathway had similar color as violacein’s. Those from CPA, predexoyviolacein, and prodeoxyviolacein producing pathways were colorless, pink, and orange, respectively. We further investigated the violacein and deoxyviolacein pathways in TX-TL. The appropriate linear DNAs were expressed in TX-TL at an equimolar amount. The products were extracted with 70% ethanol, and a UV-Vis spectrum of each was obtained (Fig. 3). The spectra demonstrated similar shapes and peak positions to those found in the literatures [5], [6].

**Fig. 2.**
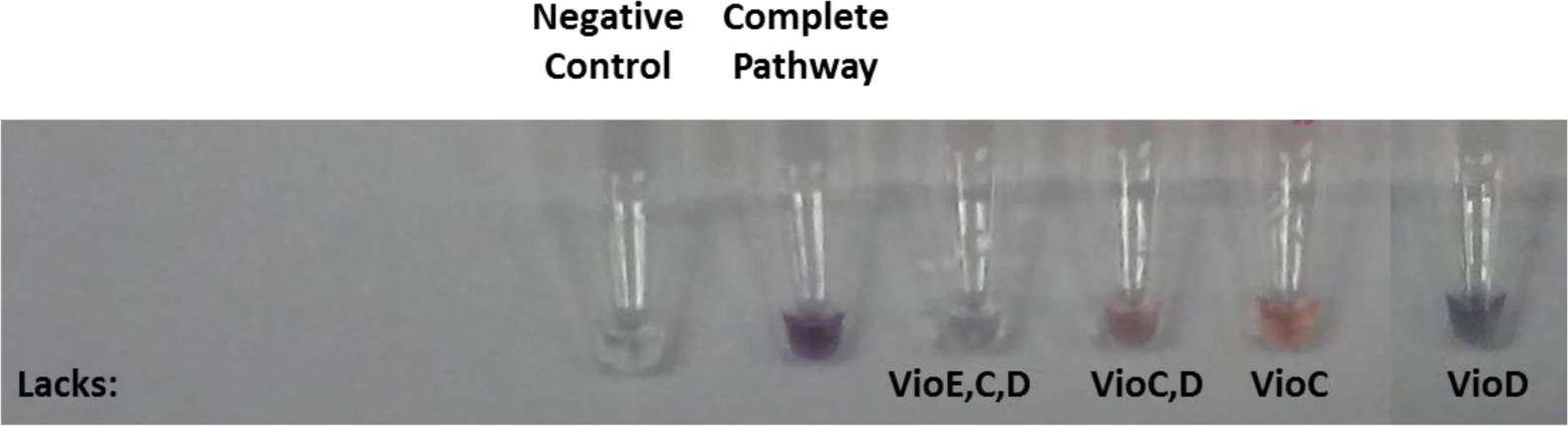
Metabolite Extraction from TX-TL Reactions. The metabolites were pelleted by centrifugation, washed with water, and re-dissolved in 70% ethanol.

**Fig. 3.**
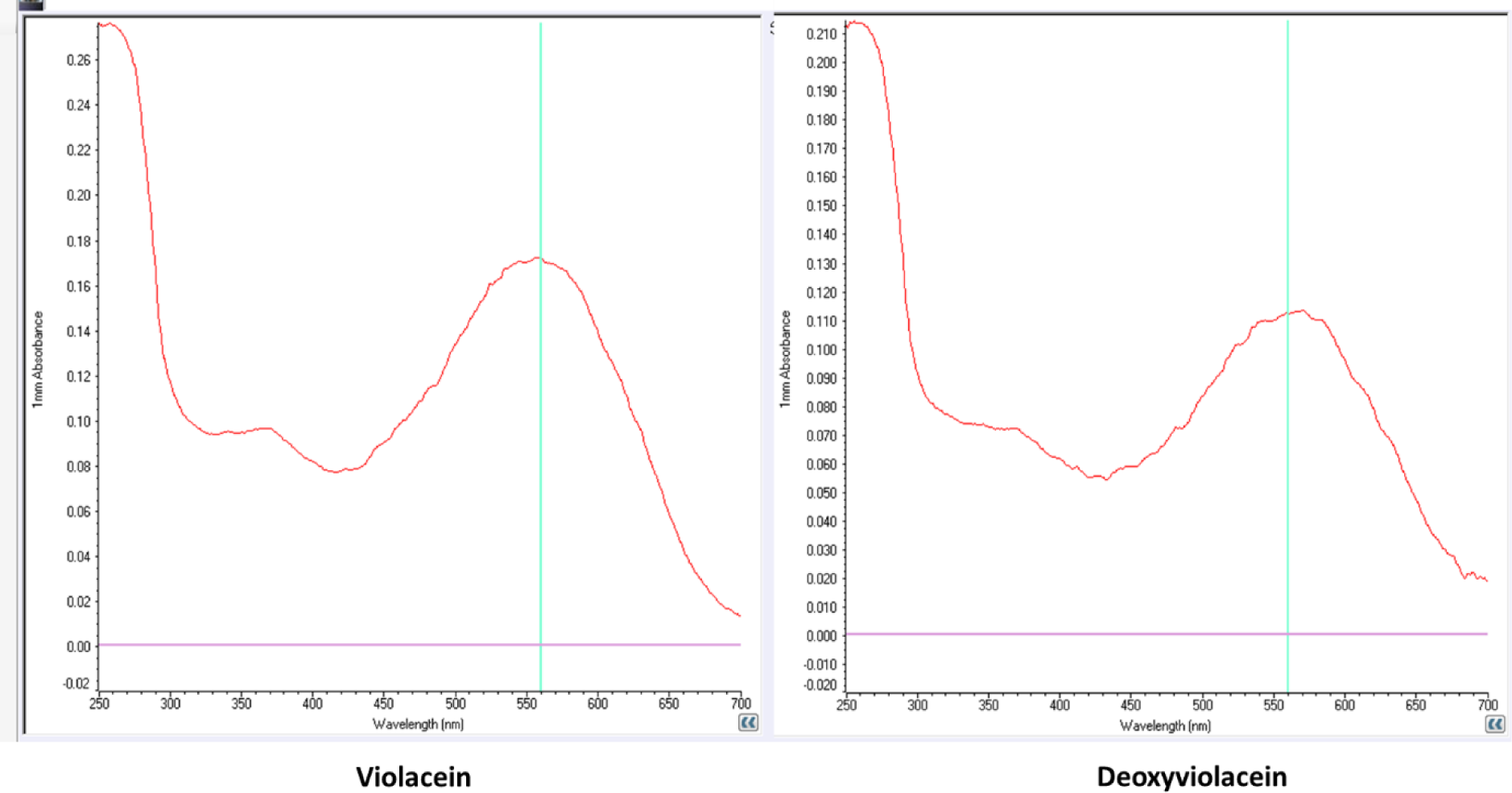
UV-Vis Spectra of TX-TL Produced Violacein and Deoxyviolacein. Metabolites from TX-TL reactions were pelleted by centrifugation, washed with water, and dissolved in 70% ethanol. The blue lines indicate the literature reported peaks [5].

### LC-MS analysis of violacein pathway reactions suggested successful synthesis of violacein in TX-TL

To further analyze TX-TL reactions of violacein pathway, products from the reactions was analyzed using LC-MS method. Commercially available violacein was used as a standard. Equimolar concentration of 4 nM for each enzyme’s DNA was used in TX-TL. Products were extracted with methanol and applied onto a reverse phase column. The metabolites were eluted with water. Mass windows of 344.103 Da, 328.108 Da, and 308.128 Da were extracted for the analysis of violacein, deoxyviolacein, and prodeoxyviolacein, respectively. Violacein was detected in reaction, confirming the functionality of the pathway in TX-TL. Based on peak area analysis, the sample contained considerable deoxyviolacein byproduct. Prodeoxyviolacein, an intermediate in the pathway, was also present in significant quantity (Fig. 4). Using a commercial standard, violacein concentration produced in TX-TL was determined to be approximately 68 ng of violacein per 10 μl reaction volume, or 19.8 μM.

**Fig. 4.**
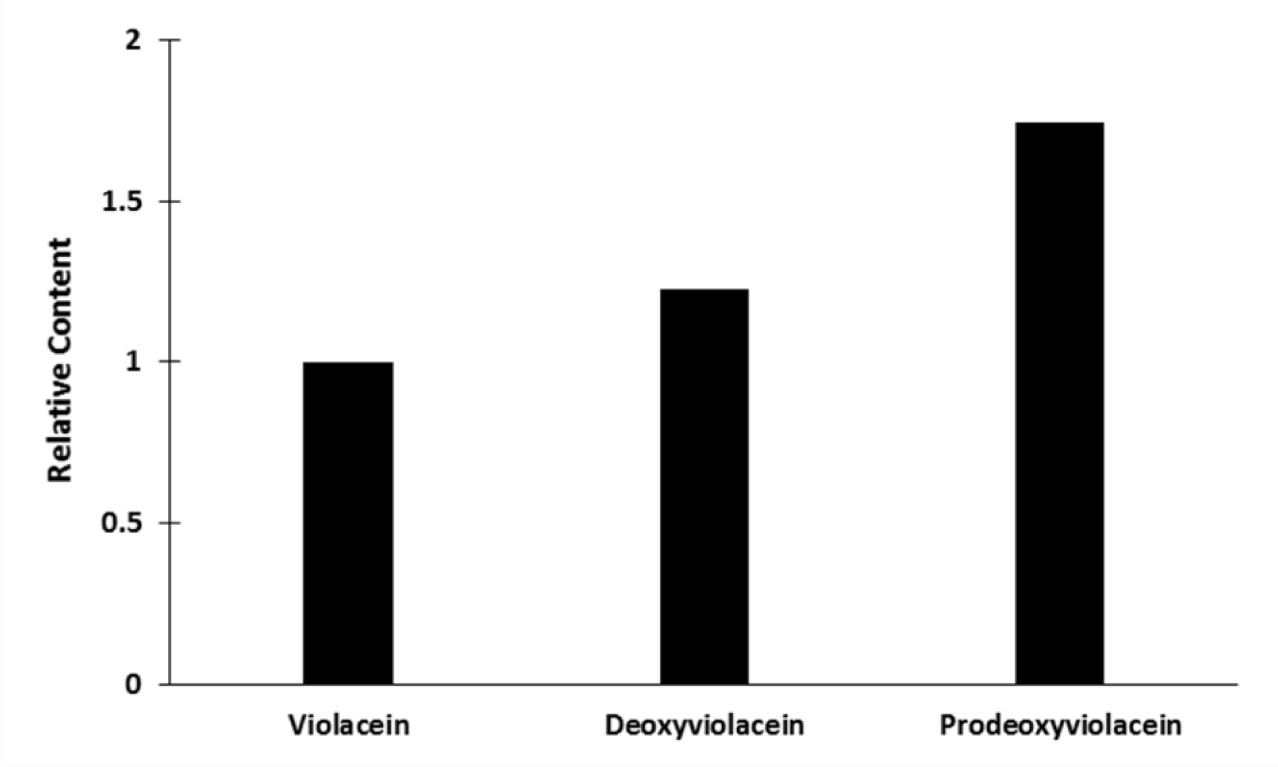
LC-MS Analysis of Violacein, Deoxyviolacein, Prodeoxyviolacein Content in 4 nM Equimolar DNA TX-TL Reaction. Peak area of each metabolite was normalized to that of violacein. LC-MS condition was the same as described in Fig. 4.

### Design space exploration of the violacein pathway in TX-TL suggested high DNA concentration of both VioC and VioD would increase violacein yield and limit intermediate buildups

To create differences in expression levels in TX-TL, various DNA concentrations of each vio enzyme were used. For each enzyme, concentrations of 2.5 nM, 4 nM, and 6.5 nM were defined as “low”, “medium”, and “high” respectively. Multiple combinations of these concentrations were added in TX-TL. The products were extracted with 70% ethanol. Crude violacein yields were determined by measuring A572 (Fig. 5). Compared to the equilmolar DNA reaction of 4 nM, an increase of VioD DNA concentration to 6.5 nM resulted in an increase in the crude yield. An additional increase of VioC DNA concentration led to a further improvement in yield. However, an increase in VioB DNA concentration instead of VioC did not improve the yield. Lowering of VioC and VioB DNA significantly affected crude violacein production.

**Fig. 5.**
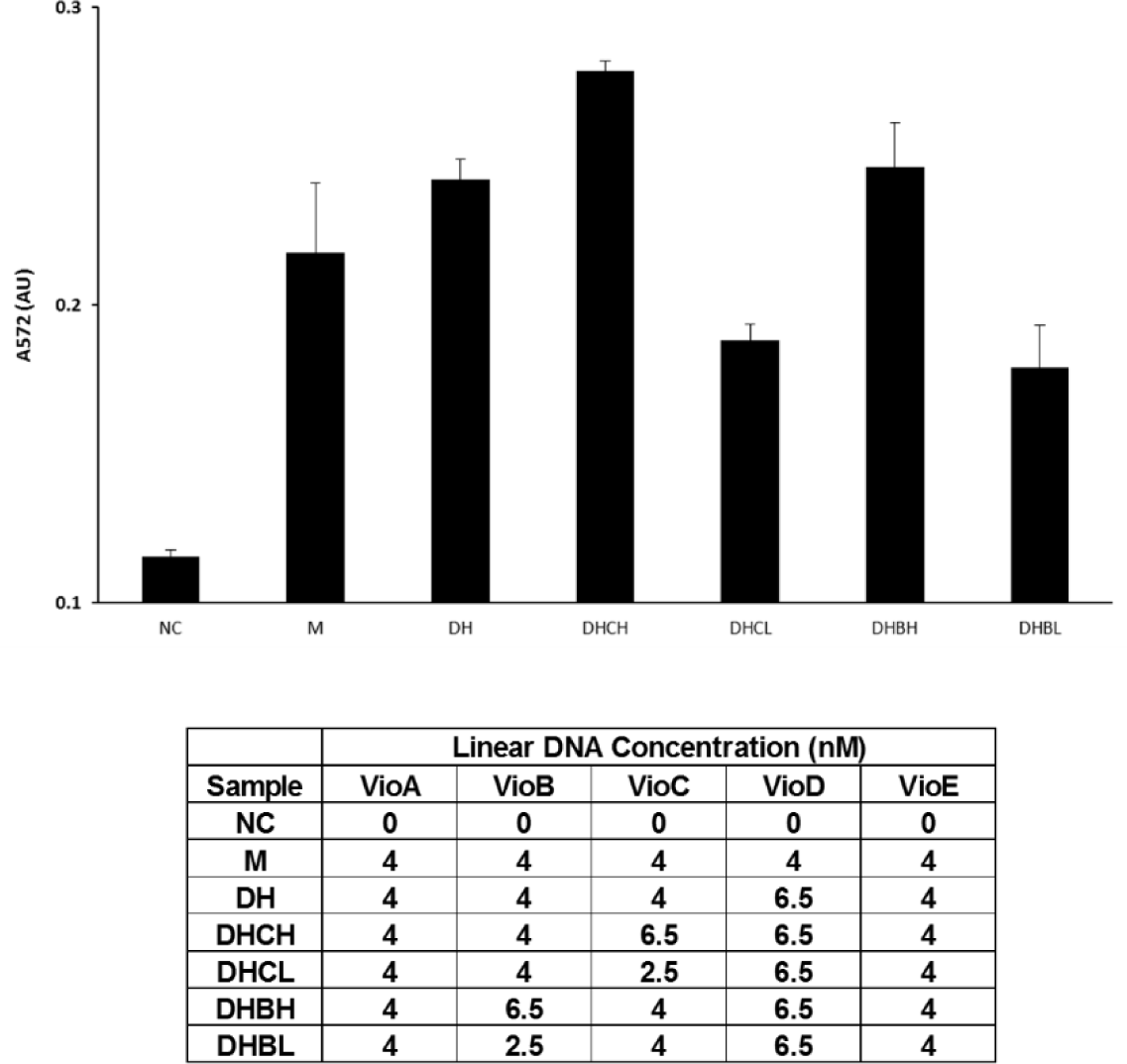
Design Space Exploration of Violacein. Metabolites from TX-TL reactions were pelleted by centrifugation, washed with water, and dissolved in 70% ethanol. Ten microliters of each sample was measured for absorbance at 572 nm for crude violacein yield estimations.

Various expression combinations of the vio enzymes were further investigated for an increase in true violacein yield. Since a buildup of prodeoxyviolacein was observed in the 4 nM equimolar DNA reaction, VioC and VioD DNA were increased to 6.5 nM to rescue the bottleneck. Additionally, TX-TL reactions with lowered VioE DNA concentration to 2 nM were ran to minimize the buildup. The products were extracted with methanol and analyzed for violacein, deoxyviolacein, and prodeoxyviolacein contents using LC-MS (Fig. 6). Reactions with 6.5 nM VioD and VioC combined with 4 nM VioE and VioB showed a high violacein content compared to deoxyviolacein and prodeoxyviolacein. Lowering VioE DNA while keeping VioB, VioC, VioD concentration high resulted in a further improve in violacein yield. However, the deoxyviolacein and prodeoxyviolacein contents also significantly increased. Other tested reactions did not show noticeable improvement in violacein production.

**Fig. 6.**
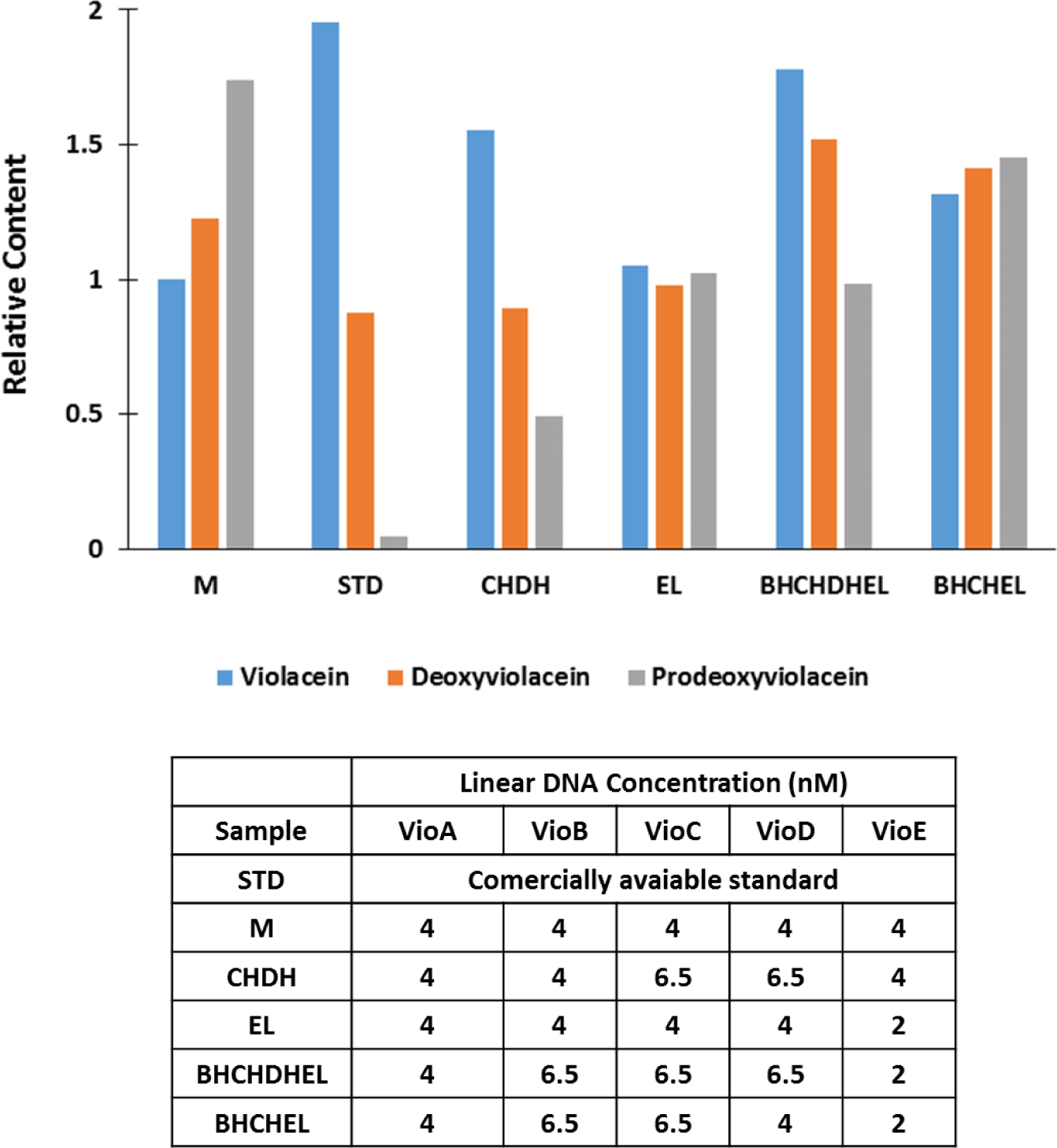
LC-MS Analysis of Violacein, Deoxyviolacein, Prodeoxyviolacein Contents in Space Exploration TX-TL Reactions. Peak area of each metabolite was normalized to that of violacein at 4 nM equimolar DNA concentration. LC-MS condition was the same as described in Fig. 4.

## Discussion

In this study, the violacein metabolic pathway has been successfully reconstructed in TX-TL. All five pathway enzymes were cloned under a medium-strong promoter and a strong RBS. The complete pathway reaction yielded product with a purple color. Omitting each of VioC, VioE, VioD, or a combination of these enzymes triggered the production of different colored metabolites. The result demonstrated the functionality of these enzymes in TX-TL. This also hints at the possible use of TX-TLto rapidly generate specific intermediates in metabolic pathways. Many intermediates are unstable, however. Therefore, preserving methods need to be investigated beforehand.

LC-MS analysis of 4 nM equimolar DNA TX-TL reaction suggested the presence of violacein at an approximate concentration of 68 ng of violacein per 10 μl reaction volume. Despite the significant buildup of prodeoxyviolacein and deoxyviolacein, this data further demonstrated the pathway’s functionality in TX-TL. To investigate the possible causes of this buildup, crude violacein production corresponding to different vioenzyme’s DNA concentration were analyzed. An increase in VioC and VioD DNA both improved the yield. However, increasing VioB DNA had no significant effect. Since VioB is upstream and both VioC and VioD are downstream of VioE, the pathway exploration result supported the LC-MS data of a prodeoxyviolacein buildup in this pathway.

Different DNA concentration combinations were sampled for an increased in violacein production in TX-TL. Reactions with VioC and VioD DNA concentrations adjusted to 6.5 nM while keeping the rest at 4 nM resulted in higher violacein yield. At the same time, both deoxyviolacein and prodeoxyviolacein contents were lowered. This is expected since an increase in VioC and VioD expressions leads to a higher consumption of prodeoxyviolacein intermediate. Assuming their expression levels were comparable at the same DNA concentration, this suggests VioD and VioC might be catalytically less efficient than VioE. The combination with high VioB, VioC, VioD at 6.5 nM and low VioE at 2 nM lead to a further increase in violacein. However, deoxyviolacein and prodeoxyviolacein productions also significantly increased. The result was unexpected since VioE DNA was two times lower than the previous combination. This might be due the increase in VioB expression which channeled the flux down the pathway’s branch point.

In conclusion, we demonstrated the feasibility of using TX-TL as a testing platform for metabolic engineering by successfully reconstructing the violacein metabolic pathway in and modulating its enzyme expression in TX-TL. Through analysis of the pathway using LC-MS, we established that high VioC and VioD DNA concentrations combined with lower, equimolar concentrations of the rest would lead to a high violacein production while minimizing unwanted intermediate and byproducts. This contributed to the overall effort of engineering a more efficient violacein production bacterial strain. Future work for this study includes *in vivo* testing of expression level combinations with highest violacein yields in TX-TL for increase in violacein production.

## Methods

### Cloning of VioA, VioB, VioC, VioD, and VioE and *in vitro* Linear DNA Assembly

Yeast plasmid harboring violacein pathway was obtained from Lee et al [2]. CDS of each enzyme was PCR amplified using New England Biolab’s Phusion® Hot Start Flex 2X Master Mix. Annealing temperature was set to 60oC for 30 sec. Elongation time was 2 minute at 72oC. The PCR was set to 35 cycles. Each CDS was ligated to its promoter, 5’ UTR, and terminator as described in Table 1 using standard Golden Gate cloning protocol described by Sarrion-Perdigones et al [7]. Complete linear DNA fragments were checked for integrity using agarose gel and sequencing.

### Preparation of crude cell (“E31”) extract

Standard *BL21-Rosetta* strain was used for extraction. Cells were grown in 1 L cultures and pelleted according to the protocol stated by Sun et al. [8] with the following modification: The 2xYT media was supplemented with 6.2% (v/v) phosphates solution (11:20 dibasic: monobasic potassium phosphate). Collected cells were homogenized and extracted according to the method described by Kwon, et al. [9] with the post-homogenization incubation period extended to 80 min instead of 60 min.

### TX-TL reactions of violacein pathway and metabolite extraction

TX-TL buffer and mastermix were prepared as described in Sun et al [8]. Extract “E31” was used in the entirety of this study. TX-TL reaction was setup by adding Mastermix to DNA mixture to the appropriate DNA concentrations. TX-TL reaction product was pelleted. The supernatant was discarded, and the pellets were washed with water once. The precipitated products from each sample were re-dissolved in 70% ethanol and measured for absorbance at 572 nm using BioTeK Synergy H1 microplate reader.

### LC-MS analysis of TX-TL produced violacein

TX-TL reaction products were pelleted and washed as described above. The metabolites were re-dissolved in methanol. The sample was injected onto Acquity UPLCr BEH C18 1.7 μm reverse phase column. Buffer A was 10 nM Ammonium Formate in 1% ACN. Buffer B was Acetonitrile. The flowrate was 0.5 ml/min. Applied gradient was from 0 to 90% B in 3 min and hold for 0.2 min. Mass windows of 344.103 Da, 328.108 Da, and 312.113 Da and elution times of 1.93, 1.65, 1.83 min were extracted for the analysis of violacein, deoxyviolacein, and prodeoxyviolacein respectively using MassLynx 4.1 software.

## Acknowledgment

We thank Michael E. Lee and the Dueber Lab from UC Berkeley for providing us the yeast plasmid with the violacein pathway. We thank Dr. Nathan Dalleska and the Environmental Analysis Center forthe support and assistance using LC/MS. We thank Dr. David A.Tirrell and Samuel Ho for guidance on the analysis of violacein. We thank Yutaka Hori and Clarmyra Hayes for the “E31” extract preparation. This work was supported by Caltech’s Student-Faculty Program and by the Gordon and Betty Moore Foundation through Grant GBMF2809 to the Caltech Programmable Molecular Technology Initiative.

